# Evolution of the Growth Hormone Gene Duplication in Passerine Birds

**DOI:** 10.1101/2022.07.29.502092

**Authors:** Shauna A. Rasband, Peri E. Bolton, Qi Fang, Philip L. F. Johnson, Michael J. Braun

## Abstract

Birds of the order Passeriformes represent the most speciose radiation of land vertebrates, yet the cause or causes of their elevated species richness have not been satisfactorily explained. One potential key adaptation is their sole lineage-specific gene, a duplicate copy of growth hormone (GH), present in all major lineages of passerines, but in no other group of birds. Growth hormone genes plausibly influence extreme life history traits that passerines exhibit, including the shortest embryo-to-fledging developmental period of any avian order. To unravel the implications of this GH duplication, we investigated the molecular evolution of the ancestral avian GH gene (GH or GH1) and the novel passerine GH paralog (GH2), using 497 gene sequences extracted from 342 genomes. Passerine GH1 and GH2 are reciprocally monophyletic, consistent with a single duplication event from a microchromosome onto a macrochromosome in a common ancestor of extant passerines. Additional chromosomal rearrangements have changed the syntenic and potential regulatory context of these genes. Both passerine GH1 and GH2 paralogs display substantially higher rates of nonsynonymous codon change than non-passerine avian GH, suggesting positive selection following duplication. A site involved in signal peptide cleavage is under selection in both paralogs. Other sites under positive selection differ between the two paralogs, but many are clustered in one region of a 3D model of the protein. Both paralogs retain key functional features and are actively but differentially expressed in two major passerine suborders. These phenomena suggest that growth hormone genes may be evolving novel adaptive roles in passerine birds.

## Introduction

Gene duplication is among the most important forces in evolution (Ohno 1970). Duplicated genes, less constrained by negative selection (Ohno 1970; Lynch and Force 2000) and in a new regulatory context, can accumulate mutations and gain new or more specialized functions (Assis and Bachtrog 2015; Jiang and Assis 2017). Duplication followed by adaptation has occurred multiple times in the vertebrate growth hormone protein family, which has expanded via whole genome and segmental gene duplications in various lineages (McKay et al. 2004; Ocampo Daza and Larhammar 2018). Here we investigate the origin, evolution, and expression of one such gene duplication that occurred at the root of the avian order Passeriformes.

Passeriformes represent a monophyletic (Raikow 1982, Hackett et al. 2008) super-radiation comprising ~6,500 of the ~10,800 extant bird species (Gill et al. 2021), making them the most speciose group of land vertebrates. The reasons for their apparent evolutionary success have long been debated. Raikow (1986) argued that the large number of passerine species is “likely an artifact of classificatory history,” prompting counterarguments proposing traits or mechanisms that may account for passerine diversity. Traits proposed include true vocal learning (Fitzpatrick 1988, Vermeij 1988) and sexually dimorphic plumage (Barraclough et al. 1995), both thought to drive speciation via sexual selection. Alternatively, passerine speciation might be driven by strictly adaptive traits, such as a generalist ability to survive in new environments (Baptista and Trail 1992), the ability to build complex, camouflaged nests (Olson 2001), enhanced neural estrogen signaling (Schlinger et al. 2022), or greater behavioral plasticity, memory, and cognition than other avian lineages (Gesicki 2019).

However, many of these proposed key adaptations are not restricted to passerines. Vocal learning is present in parrots and hummingbirds (Johnson and Clark 2020). Sexually dimorphic plumage, found in almost every avian order, is likely the ancestral state for birds (Kimball et al. 1999). Complex nest building is not unique to passerines either; Olson (2001) cites the example of the mud nests built by swallows without acknowledging the equally complex nests made by swifts and other birds (Steeves et al. 2020). The cognitive abilities of parrots are also highly developed (Emery 2006). Biogeographical explanations of passerine species diversity are likewise inadequate; passerine diversification rates, while linked to major climatological and geological events, are decoupled from continent colonization events (Oliveros et al. 2019), and, while island systems have been shown to promote diversification of some passerine clades (McCullough et al. 2022), it is not clear why this effect would not drive commensurate net diversification in other avian orders. Thus, after over 40 years of active consideration, there are no clearly favored hypotheses on the causes of passerine diversity.

What then remain as possible mechanisms or key adaptations that might explain passerine diversity? Whether high speciation rates, low extinction rates, or both have driven passerine diversity since the origin of the clade ~47 Ma (Oliveros et al. 2019), there may be underlying adaptive mechanisms. Compared to other birds, passerines have rapid developmental periods, short generation times, and many offspring (Cooney et al. 2020). Such traits are likely influenced by hormonal physiology. Indeed, hormones of the growth hormone/insulin-like growth factor axis are known to influence several aspects of development and life history in birds such as body size, time to reach maturity, clutch size, egg weight, clutch interval, and lifespan (Lodjak et al 2018; Socha and Hrabia 2019). Could adaptive innovations in hormone physiology contribute to the evolutionary success of passerines?

Growth hormone (GH), also called somatotropin, is the ancestral hormone of the GH protein family, controlling adult body size across all vertebrates (Kawauchi et al. 2002). The expansion of the growth hormone family via gene duplication in other taxa has led to a variety of functional hormones with novel adaptive functions, such as the convergent evolution of placental lactogens in disparate eutherian mammalian lineages (Papper et al. 2009; Alam et al. 2006). GH family genes have also evolved distinct and overlapping functions in diverse lineages. For example, prolactin, ancestrally an osmotic regulator in fishes, (Ocampo Daza and Larhammar 2018), drives milk secretion in mammals (Soares 2004), crop milk production in pigeons and doves (Horseman and Buntin 1995) and brooding behavior in diverse birds (Angelier 2009).

Yuri et al. (2008) described a duplication of the GH gene in passerine birds and presented evidence for accelerated evolution and positive selection of the paralogs. Arai and Iigo (2010) independently discovered this duplication and provided evidence of differential expression of the two genes. With the recent release of hundreds of newly assembled genomes from virtually every family of extant birds, a search for taxon-limited genes in Passeriformes found that the growth hormone gene duplication is the only novel gene or gene copy universally present in this clade (Feng et al. 2020), spotlighting the possibility that this genomic innovation could be correlated with passerine diversification.

Here, we evaluate molecular evolution of the growth hormone paralogs using complete GH sequences from 183 passerines and 159 other birds. The rich genomic resources now available allowed us to examine the chromosomal context of both paralogs in multiple species and resolve the phylogenetic placement of key events including the GH duplication, lineage-specific deletions, and rearrangements of syntenic genes. In addition, we provide the first RNA-seq evidence on transcriptional functionality and tissue specificity of both GH paralogs in a passerine and speculate on the potential neofunctionalization or subfunctionalization of these genes in the major lineages of passerines.

## Results

### Copy number and duplication origin

We retrieved a total of 497 GH genes from 342 bird genomes including all major lineages of living birds, comprising 159 non-passerines with a single copy, 154 passerines with two paralogs retrieved, and 29 passerines with only one paralog retrieved (Supplemental Table 1). BLAST searches did not identify more than one copy in any non-passerine or more than two copies in any passerine. Gene tree estimation demonstrates that passerine GH sequences form reciprocally monophyletic clades that split at the root of Passeriformes (Figure 1), consistent with a single GH gene duplication event in living birds. We designate the paralogs belonging to these clades GH1 and GH2 (see Methods for naming rationale). All taxa with two paralogs had one in each clade, and taxa for which we recovered only one paralog (eight with GH1, 21 with GH2) were widely dispersed on the tree (Figure 1). Lack of a clear phylogenetic pattern to these absences suggests that most or all missing paralogs are due to the draft or low-coverage nature of many bird genomes, as previously concluded by Feng et al. (2020).

**Figure 1:**
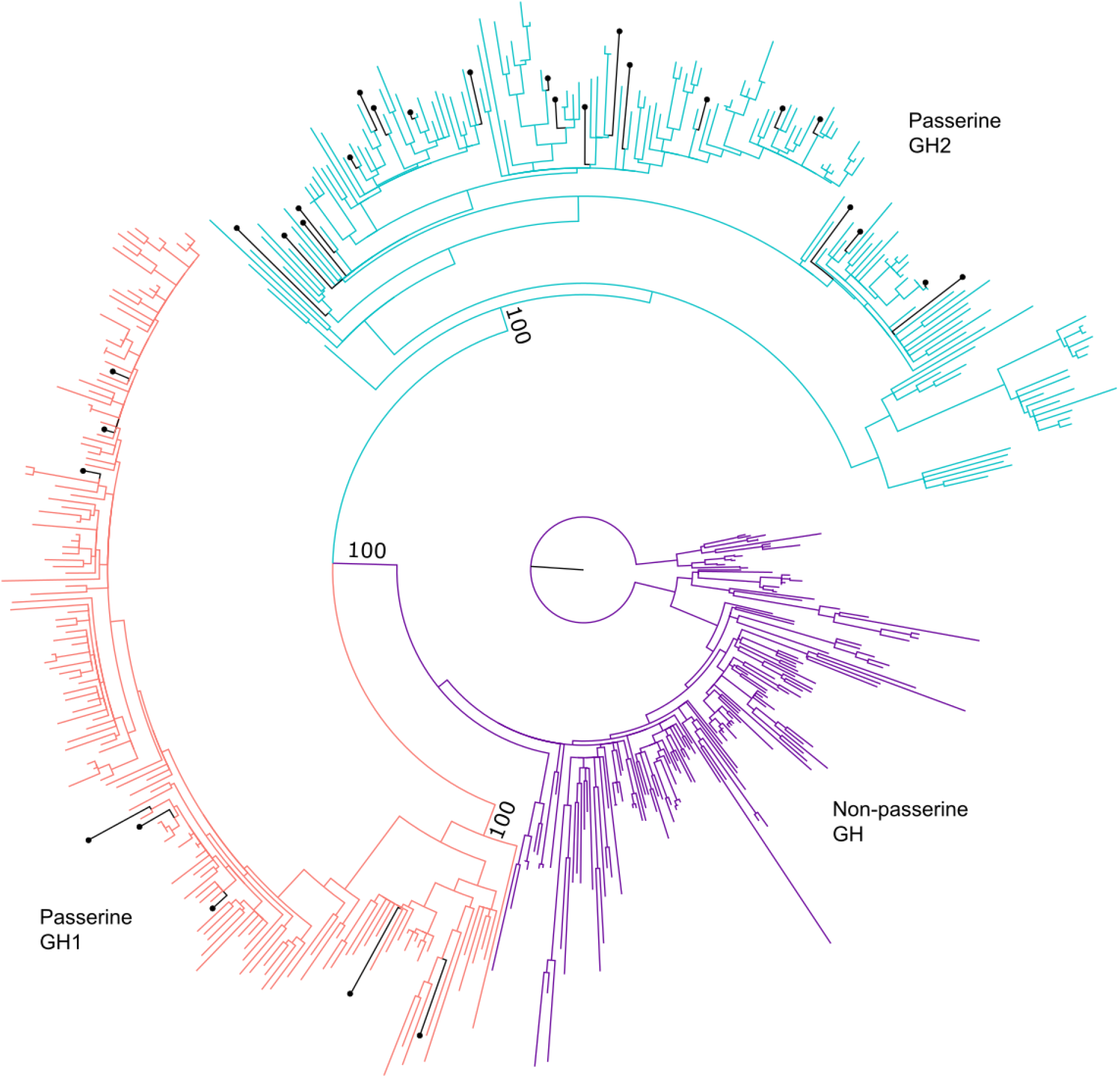
Maximum likelihood gene tree of avian growth hormone (GH) genes. The tree search was based on the nucleotide sequences of GH coding regions and was constrained to topologies compatible with widely accepted species relationships of birds well supported by phylogenomic studies (Supplemental Fig. 2). Major clades are labeled and colored: non-passerine GH in purple, passerine GH1 in salmon, and passerine GH2 in teal. Bootstrap values supporting reciprocal monophyly of passerine GH1 and GH2 clades are shown, identical to the ones obtained in the unconstrained topology. Passerine species with a single paralog retrieved have black branches marked with circle tips.

The syntenic context of avian GH genes lends further evidence to a single GH gene duplication event and shows that this event is part of a series of genic and genomic changes. Of 52 non-passerine genomes in which we examined syntenic context, GH was adjacent to CD79B in 46 genomes and the remaining 6 genomes had incomplete assemblies, where GH was the sole gene annotated on a scaffold not much longer than the gene itself. Similarly, in 24 passerine genomes in which we examined syntenic context, the GH1 gene was also adjacent to CD79B in all cases. This linkage group resides on microchromosome 27 of chicken (*Gallus gallus*) and zebra finch (*Taeniopygia guttata*). As the linkage of CD79B and GH occurs across Tetrapoda, e.g., *Xenopus tropicalis* genome assembly UCB_Xtro_10.0 and *Homo sapiens* genome assembly GRCh38.p13 (O’Leary 2016), we infer this syntenic context to be ancestral and passerine GH1 to be orthologous to the single-copy non-passerine GH (Figure 2).

**Figure 2:**
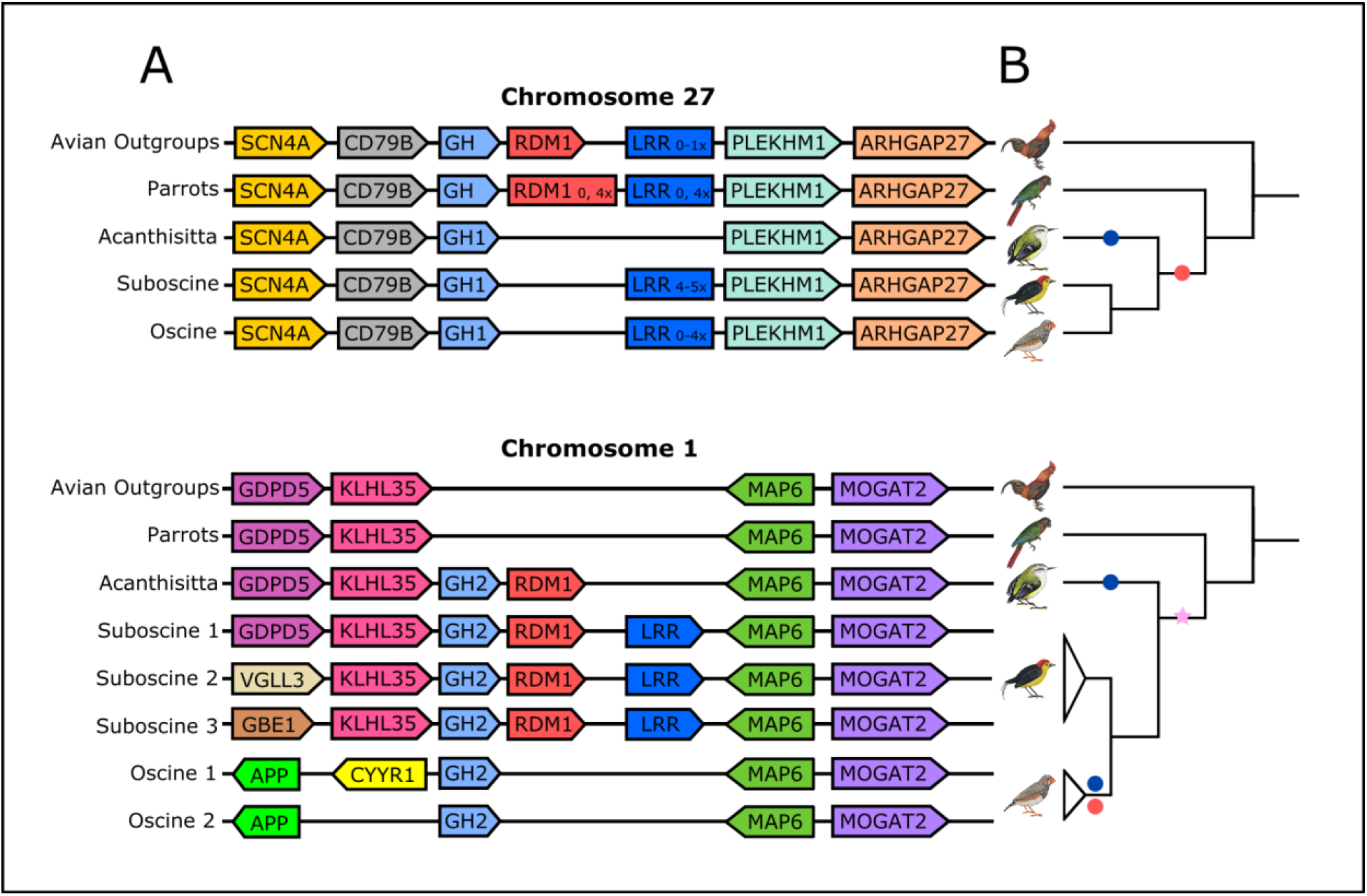
***A-** Local syntenic contexts of GH genes on chromosome 27 (ancestral locus) and chromosome 1 (duplicate locus in passerines). Colored pentagons indicate relative gene placements and orientations (distance between genes not to scale). Rectangles indicate a variable number of tandem copies of that gene. Post-duplication, many changes to local syntenies have occurred in passerines. Each row in the diagram represents a local synteny pattern found in multiple species in avian outgroups, the sister taxon to passerines (parrots), and each passerine suborder, except for* Acanthisitta, *for which only one genome exists. The underlying syntenies informing this diagram, as well as a few single species exceptions to it, are reported in Supplemental Table 8*. **B-** Cladograms of relationships among the passerine suborders and outgroups depicted in part A, with symbols denoting key rearrangements on chromosomes 27 and 1. **Star:** segmental duplication of a part of chromosome 27 containing GH, RDM1, and an LRR (leucine-rich repeat) gene onto chromosome 1. **Navy circles:** deletions of LRR genes. **Red circles:** deletions of RDM1 genes.

In this same set of 24 passerine genomes, the derived paralog (GH2) consistently occurs near MAP6 in a syntenic context that maps to zebra finch chromosome 1 (Figure 2). This pattern is consistent with an interchromosomal duplication of a ~20 kb segment containing GH1 from ancestral chromosome 27 onto ancestral chromosome 1, with a breakpoint only 4 kb upstream of the GH translation start site (Supplemental Figure 1). Further rearrangements on chromosome 1 have changed the syntenic context immediately upstream of GH2 in oscines (suborder Passeri), potentially affecting the regulatory context of this paralog (Figure 2; Supplemental Figure 1). The ancestral segment containing GH that was duplicated from chromosome 27 to chromosome 1 also included two other predicted genes downstream: RDM1 and an unnamed leucine-rich repeat (LRR) gene. A series of deletions on chromosome 1 and tandem duplications or deletions on chromosome 27 have changed the number of RDM1 and/or LRR copies in various lineages, while GH1 and GH2 are widely, possibly universally, retained in passerines (Figures 1&2).

### Gene structure

Passerine GH1 and GH2 share the same gene structure, with five exons and four introns. Like human GH1, avian GH genes are typically 217 amino acids in length (observed range 214-218 residues). Small indels of 1-3 codons in single taxa or clades account for the length variation (Supplemental Table 2).

Like those of other exported proteins, GH amino acid sequences typically begin with signal peptides that target them for secretion. The vast majority of non-passerine GH sequences are predicted with high likelihood to have signal peptides: 158 of 160 species examined across 38 orders (Supplemental Table 3). Most passerine GH1 sequences are also predicted with high likelihood to have signal peptides, including 156 of 162 sequences examined. In contrast, many fewer passerine GH2 sequences are predicted to include a signal peptide (116 of 175 examined), and the likelihoods of those that do are often substantially lower. The predicted lengths of signal peptides range from 24 to 29 residues, with 27 residues being by far the most common (Supplemental Table 3).

Untranslated regions (UTRs) annotated in passerine GH paralogs were somewhat longer in GH2 than in GH1 (Supplemental Table 4). For 5’ UTRs, GH1 had a mean length of 54.47 (range 21-97) bp, while for GH2, the mean was 70.24 (range 59-101) bp. For 3’ UTRs, GH1 had a mean of 48.93 (range 22-71) bp, versus a mean of 85.74 (range 83-100) bp for GH2. UTR lengths are more variable in the much broader taxonomic distribution of annotated non-passerine avian GH paralogs, which have 5’ UTRs from 28-119 bp (mean 56.5) and 3’ UTRs from 18-112 bp (mean 86.33).

The base composition of the passerine paralogs has diverged post-duplication (Figure 3). While the GC content of both paralogs must initially have been identical, all three passerine suborders now display substantial differences between GH1 and GH2 in GC content (mean difference = 4.87; range 1.42-8.12 across all species; Supplemental Table 5). However, GH1 has higher GC content in *Acanthsitta* (suborder Acanthisitti) and oscines, while GH2 is higher in suboscines (suborder Tyranni).

**Figure 3:**
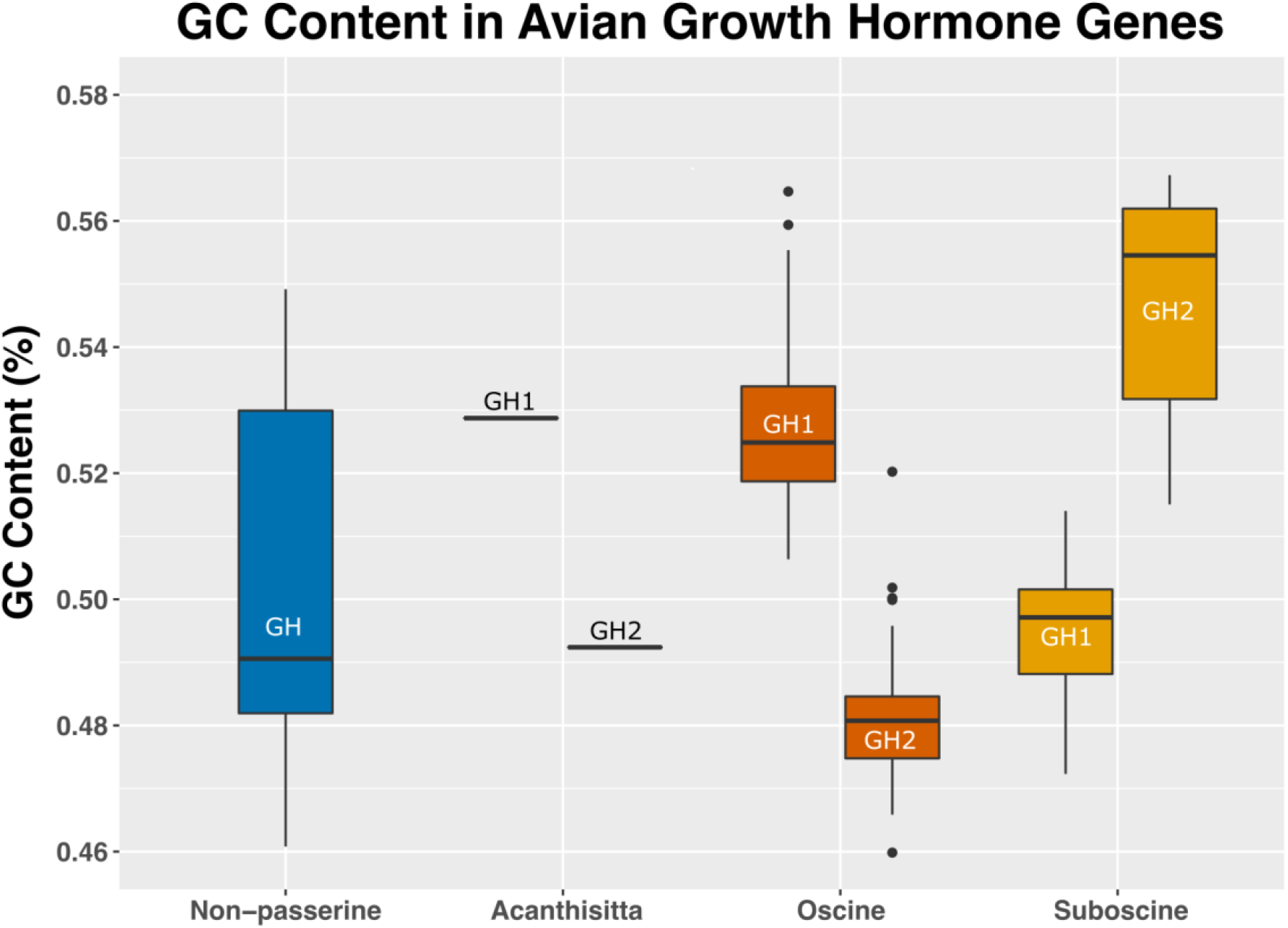
Base composition (G+C percentage) of avian GH genes (exons plus introns) by group and paralog. Non-passerine GHs are represented by 24 species in 16 orders. Acanthisittines are represented by one species with one copy of each paralog, whereas the other suborders are represented as follows: suboscines- 22 GH1, 21 GH2; oscines- 97 GH1, 105 GH2. See Supplemental Table 5 for the base composition per species and paralog.

### Accelerated evolution and positive selection in the passerine growth hormone paralogs

Visual examination of the GH gene tree based on protein-coding sequences suggests that passerine GH paralogs generally have longer branches than non-passerine GH (Figure 1). We confirmed this pattern with a likelihood ratio test using baseml (paml v. 4.9j; Yang 2007), allowing for three different molecular clock rates: a background rate, a rate for the passerine GH1 clade, and a rate for the passerine GH2 clade. This test strongly rejected all three rates being identical (p <0.00001; Supplemental Table 6A) in favor of increases for both GH1 and GH2. These results were robust to use of a gene tree estimated strictly from the GH data (“unconstrained tree”) as well as a gene tree informed by well-accepted avian relationships (“constrained tree”; see Methods and Supplemental Figure 2). Both paralogs experienced similar post-duplication rate increases with maximum likelihood estimates of 4.8x/5.9x (constrained/unconstrained topology of the input tree) for GH1, and 4.3x/5.5x for GH2.

Since the increase in clock rate could have arisen due to neutral processes, we next conducted formal tests of positive selection. Using codeml (paml v. 4.9j; Yang 2007), we tested a null model where the ratios of nonsynonymous to synonymous substitution rates (dN/dS or ω) must be less than one, simulating no positive selection, versus three alternate models where sites with ω > 1 were allowed on foreground branches. The best-fitting model tested (likelihood ratio test against the null p < 0.0001) allowed positive selection on both passerine GH1 and GH2. Of the single-clade models, both passerine GH1 and GH2 were individually highly significant for the model with ω > 1 allowed, but the model with positive selection allowed on GH2 displayed a larger improvement (Supplemental Table 6B). Similar results were obtained for both constrained and unconstrained tree topologies.

Next, we performed a more conservative aBSREL test of positive selection (Smith et al. 2015) on the two ancestral branches leading to the passerine GH1 and GH2 clades. Significant positive selection was indicated only on the branch leading to the GH2 clade (p = 0.0429 with the constrained topology, p = 0.0622 with the unconstrained topology, both after Holm-Bonferroni correction for multiple testing). This result suggests a burst of positive selection immediately following the duplication event. Overall, the relative rate and selection test results provide evidence for positive selection on both passerine GH1 and GH2, with a stronger signal on the GH2 clade.

The Bayes Empirical Bayes test (paml v. 4.9j; Yang 2007) identified 11 amino acid sites likely under positive selection in GH1 and 11 amino acid sites likely under positive selection in GH2, with one of those sites, passerine amino acid 27, identified in both paralogs (Figure 4, Supplemental Table 6C). Position 27 is located −1 to the signal peptide cleavage site; cleavage is predicted to occur between amino acids 27 and 28 in the majority of avian GHs (Supplemental Table 3). One site identified in GH2 as likely under positive selection, site 99 as labeled herein, matches one of two such sites identified by Yuri et al. (2008) with partial sequence data. Amino acids under positive selection are clustered in the coding sequence, with the densest cluster of sites spanning the end of exon 4 and beginning of exon 5, generally avoiding areas where alpha helix structures were predicted (Figure 4).

**Figure 4.**
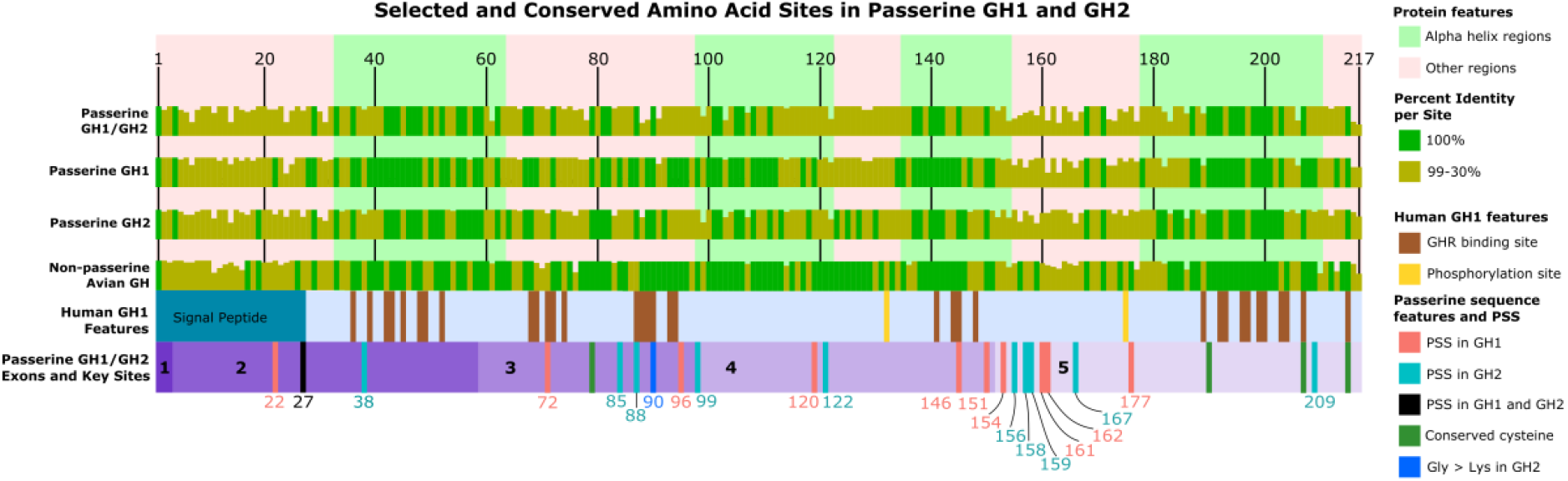
***from top to bottom:** Protein features (alpha helices and non-alpha helix regions) predicted from* Taeniopygia guttata GH1 *using AlphaFold v. 2.1.1 (Jumper et al. 2021, Varadi et al. 2021) are marked along the 217 amino acid avian consensus sequence. The percent identity at each amino acid site of various groups is overlaid, with 100% conserved sites marked in green. Insertions with less than 10% frequency are masked. Key features of human GH1 are shown aligned with avian GH sequences, including the signal peptide, sites that bind growth hormone receptor (GHR), and phosphorylation sites. At the bottom, the passerine GH1/GH2 consensus sequence bar with numbered exons shows the placement of positively selected sites (PSS). Only sites with p > 0.95 according to the Bayes Empirical Bayes test of PAML v. 4.9j (Yang et al. 2007) are shown, labeled by amino acid position and colored according to paralog. Four cysteine sites, universally conserved in avian and human GH as well as other GH family proteins (Ocampo Daza and Larhammar 2018), are marked in the passerine sequence, as is site 90, which is fixed for glycine in all GH1 and non-passerine GH sequences, but fixed for lysine in all GH2 sequences*.

### Sequence conservation and structural implications

Other regions of the proteins show varying levels of sequence conservation across birds (Figure 4). Non-passerine GH is more conserved than passerine GH1 and GH2, despite spanning a much longer period of evolutionary divergence: ~47 MY for passerines (Oliveros et al. 2019), 90-100 MY for extant lineages of non-passerine birds (Jarvis et al. 2014; Kimball et al. 2019). Overall amino acid sequence divergence between GH1 and GH2 within species varies from 5.07% to 17.51%, with a mean divergence of 13.44% for oscines and 10.04% for suboscines (Supplemental Table 7). While many sites are variable across the alignment, there is only one fixed difference between the paralogs studied here; site 90 is always glycine in GH1 and always lysine in GH2. Site 90 is also glycine in all non-passerine GH studied, thus lysine in GH2 is the derived condition at this site. This site is involved with growth hormone receptor binding in human GH1 so the change from a small, nonpolar side chain (glycine) to a long, charged one (lysine) may be of functional significance (Figure 4).

Other features of interest include two pairs of cysteines known to form two disulfide bridges in GH (Ocampo Daza and Larhammar 2018). These are universally conserved. Four predicted alpha helices are generally more conserved than the rest of the polypeptide. Sites crucial to binding with the growth hormone receptor (GHR) in human GH1 (Lu et al. 2020) are generally not under positive selection in passerine GH1 or GH2 (Figure 4), in contrast to mammalian GH paralog evolution, where positively selected sites tend to cluster in these GHR-binding areas (Buggiotti and Primmer 2006). Serine site 133 can be phosphorylated in human GH1 (Giorgianni et al. 2004), and is widely conserved in all birds (Figure 4). Site 175 can also be phosphorylated in human GH1, but not in birds. However, nearby site 177 is a serine or threonine in most bird lineages and thus potentially phosphorylatable (Supplemental Data: GH Amino Acid Alignment). This site is under positive selection in passerine GH1, where Ser has frequently been replaced by amino acids (Gly, Ala, Arg, Asn) that cannot be phosphorylated.

The potential importance of both conserved features and positively selected sites to the tertiary structure of the protein is evident in a 3D model of the folded polypeptide (Figure 5). The four conserved alpha helices form a structural core, a shared feature of GH family proteins (Ocampo Daza and Larhammar 2018). The four conserved cysteine residues form two closely apposed pairs, as needed for two disulfide bridges. Many sites aligning with the sites that in human GH1 bind to the human GHR are clustered in three of the core alpha helices and two smaller helices that are all accessible on the surface of the 3D model (Figure 5B). One core alpha helix that is buried in the model has no such sites. Both potentially phosphorylatable sites also appear on the surface of the model (Figure 5B). Finally, 13 of the 21 sites inferred to be under positive selection, as well as the fixed Gly>Lys at site 90 in GH2, are clustered near one end of the structural model (Figure 5A), although these sites are widely spaced in the primary sequence, spanning residues 85-209. They include sites under positive selection in both paralogs (six in GH1, seven in GH2) and one site (209) that lies near the end of a highly conserved region of the primary sequence that is bent into a surface loop by a disulfide bridge that lies near the carboxy terminus of the polypeptide (Figure 5B).

**Figure 5:**
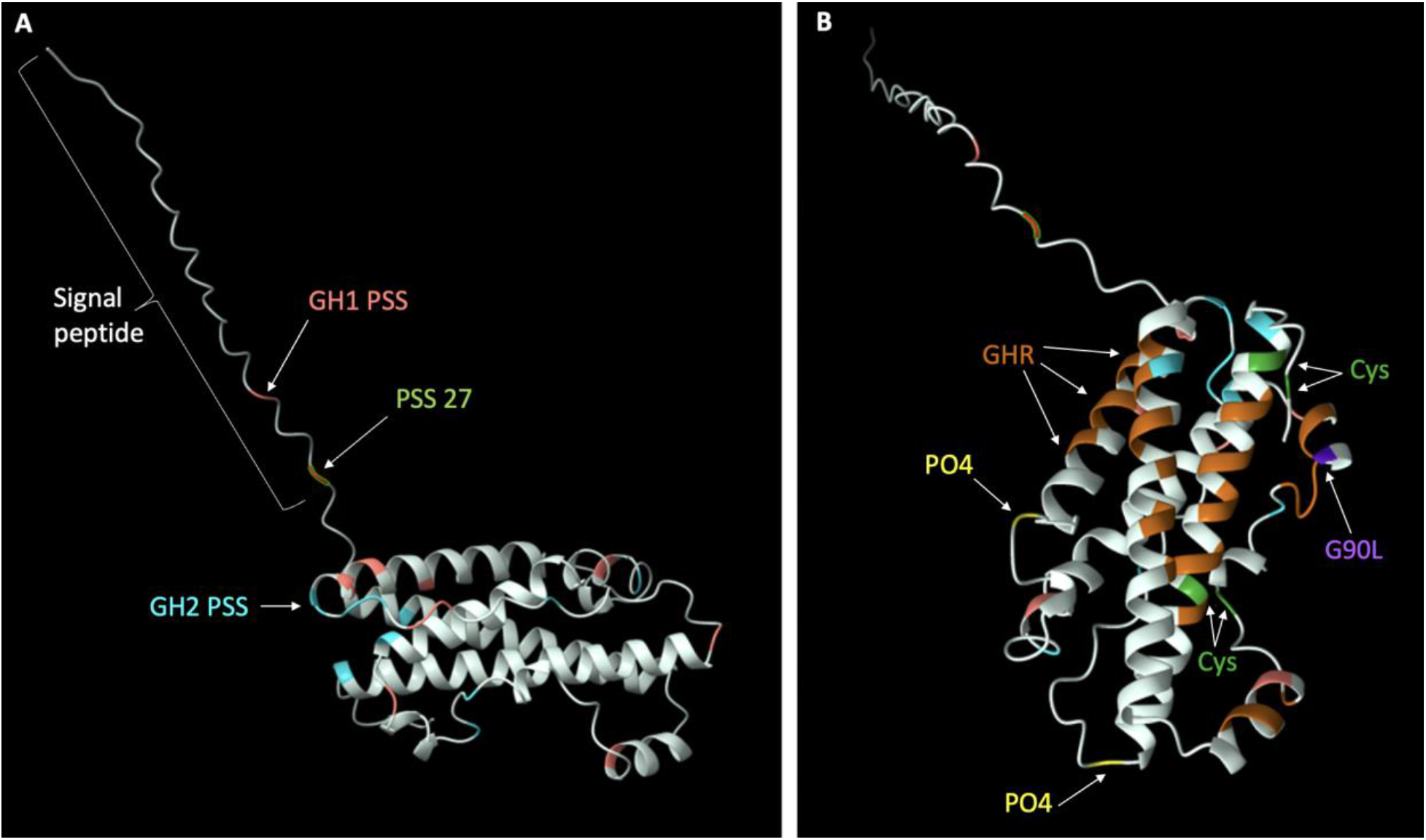
*Model of the chicken* (Gallus gallus) *GH1 tertiary structure calculated by AlphaFold v. 2.1.1. **A:** The signal peptide and exemplar positively selected sites (PSS) are labeled. All PSS are colored; those in GH1 in salmon, those in GH2 in blue and those in both in red with green outline. **B:** PSS colored as in A. Cys: conserved cysteines. PO4: potential phosphorylation sites. GHR: sites aligning with ones known to bind the human GH receptor are colored in brown. G90L: Site 90 is glycine in all GH1 and lysine in all GH2 studied here*.

### Changes to non-coding regions

The promoter region of chicken GH has been well-characterized (Ip et al. 2004). In this region, the proximal TATA box is retained in both passerine GH paralogs (Figure 6). The binding site for transcription factor Pit-1 is also likely retained, but with some sequence changes in the first few bases. The core binding site for the inhibitory thyroid hormone response element (TRE) is likewise conserved. Interestingly, a second TATA box upstream shows paralog-specific conservation and degeneration. The ancestral sequence (TATAAAT) is conserved in all passerine lineages except GH1 of *Acanthisitta* and both GH1 and GH2 of suboscines. While the function of this putative TATA box is unknown, multiple TATA boxes in promoter regions are associated with multiple transcription initiation sites in other eukaryotic genes (Hahn et al. 1985; Grace et al. 2004). All passerine GH introns appear to retain splice site functionality with the preservation of GU-AG sites, except in intron 1 from the suboscine *Tachuris rubigastrus* GH1 (data not shown).

**Figure 6:**
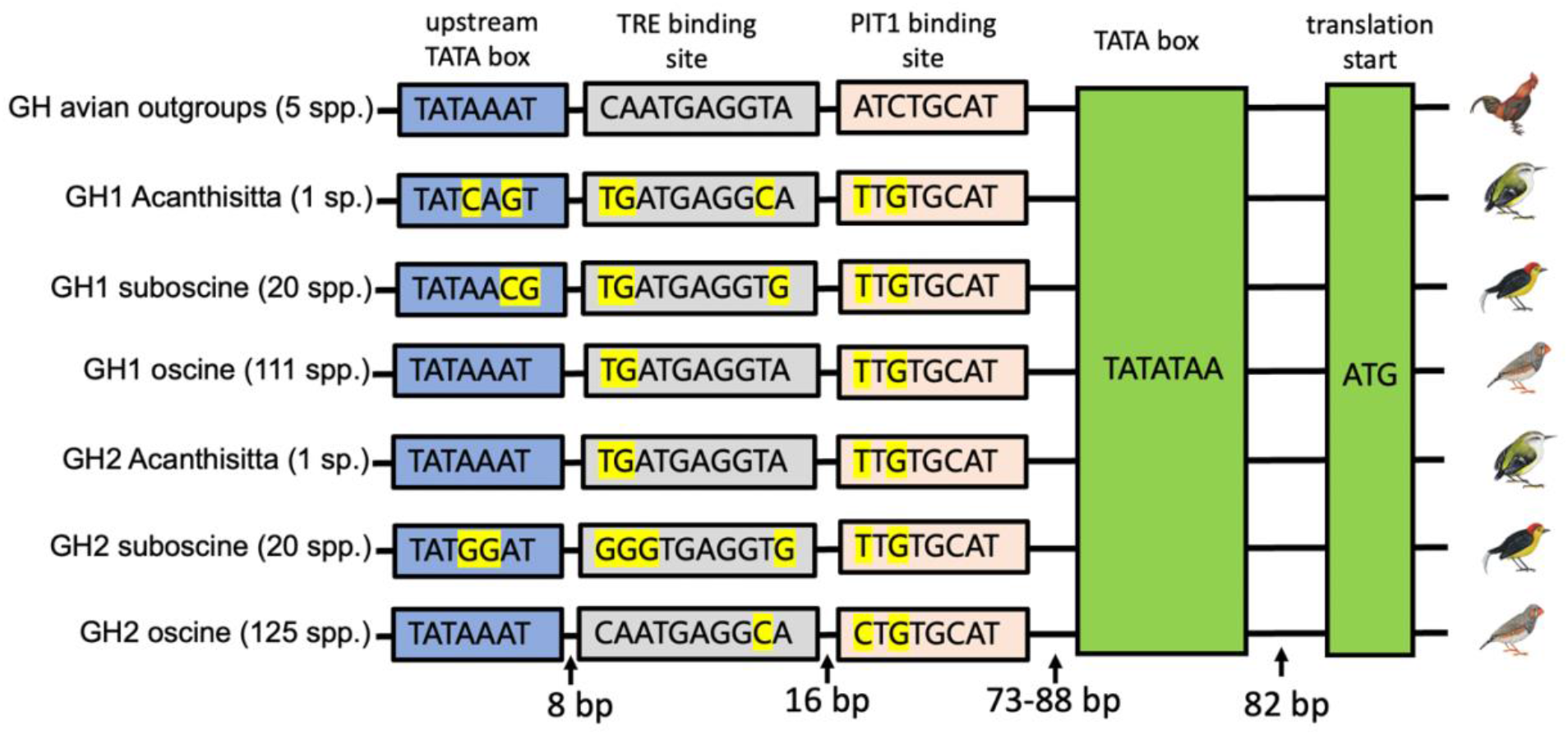
*Consensus sequences of cis-regulatory elements upstream of the translation start site in GH genes of various passerine lineages and non-passerine outgroups. Pit1(pituitary-specific transcription factor 1) is the primary transcription factor regulating expression of GH paralogs in the pituitary (Harvey et al. 2000, Mukherjee and Porter 2012). TRE (thyroid response element) is an allosteric inhibitor of the Pit1 binding site. Spacing between the PIT1 binding site and the canonical TATA box is variable across species due to insertion of a 15 bp direct repeat early in the oscine radiation that was repeatedly lost in various sublineages (Supplemental Figure 3). Non-passerine outgroups used were* Apteryx australis, Nothoprocta perdicaria, Gallus gallus, Anas platyrhynchos and Cuculus canorus.

### Gene expression

Transcriptome sequencing from multiple tissues provides the first direct evidence for the expression of both GH1 and GH2 in a suboscine, the wire-tailed manakin (*Pipra filicauda*). GH1 was expressed at a high level in the pituitary gland, at moderate levels in testes and at lower levels in ten brain areas tested (Figure 7). Brain area expression of GH1 was variable between individuals, but, after Benjamini-Hochberg correction for multiple testing, seven of ten brain regions tested still showed significant expression. In contrast, GH2 expression was detected only in pituitary at moderate levels (Figure 7). These expression patterns in our suboscine data differ from those previously reported in oscines (Arai and Iigo 2010; Xie et al. 2010) (see Discussion).

**Figure 7:**
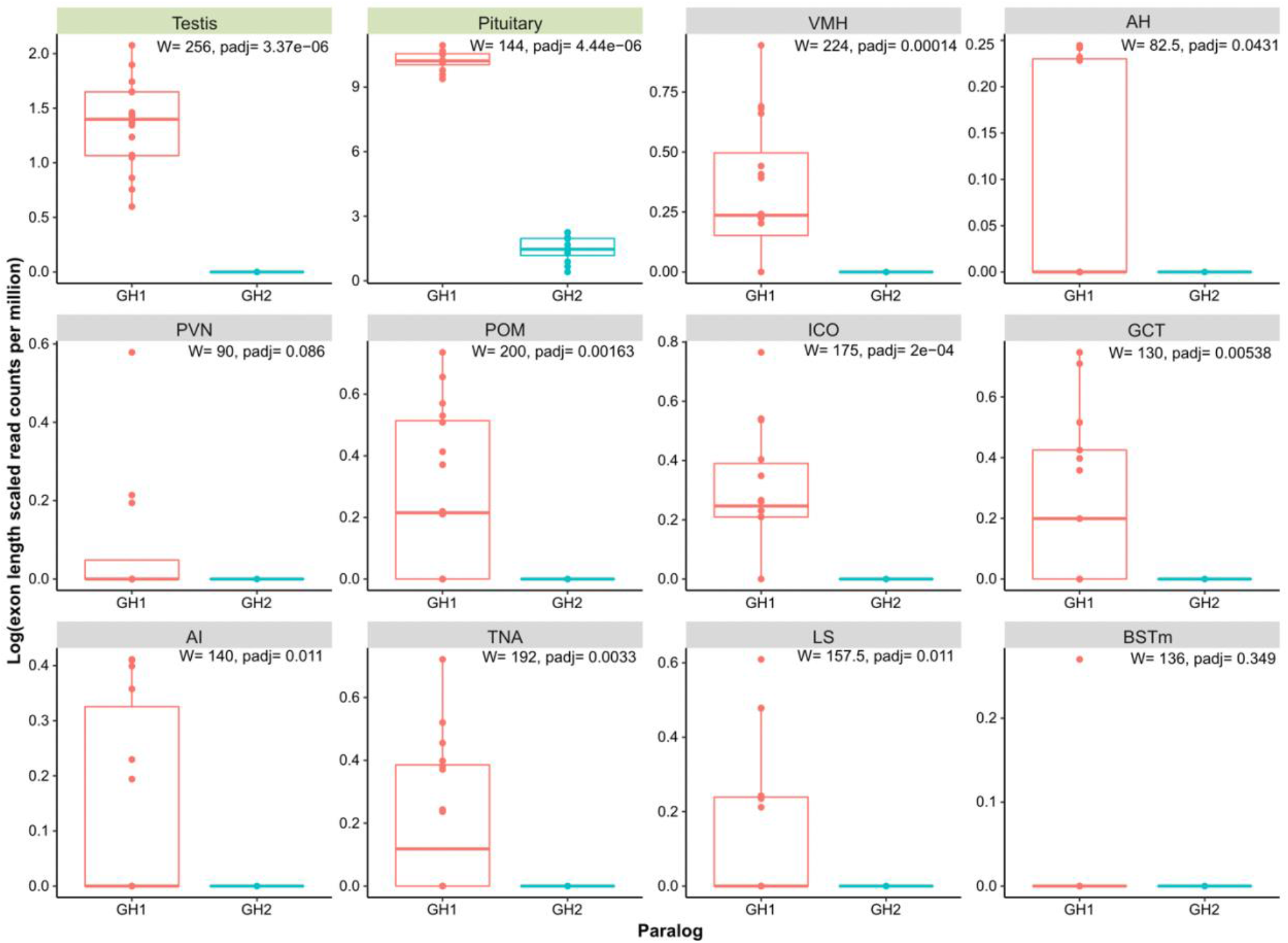
*Gene expression of GH1 and GH2 as measured by transcriptome sequencing of mRNA from tissues from 12 male* Pipra filicauda *individuals. Peripheral tissues (green heading): testis, pituitary. Brain tissues (grey heading): ventromedial hypothalamus (VMH), anterior hypothalamus (AH), paraventricular nucleus (PVN), medial preoptic area (POM), dorsomedial intercollicular nucleus (ICO), midbrain central gray (GCT), arcopallium intermedium (AI), nucleus taenia (TNA), lateral septum (LS), bed nucleus of the stria terminalis (BSTm). W = Wilcoxon test statistic of the differential expression between the two paralogs, per tissue. padj = p-value adjusted for multiple testing (Benjamini-Hochberg*).

## Discussion

Multiple lines of evidence now combine to strongly suggest a role for selection in passerine GH paralog evolution. First, both growth hormone paralogs are apparently retained in all passerines despite at least four post-duplication deletion events affecting other genes included in the segment originally duplicated (Figure 2). Second, formal tests analyzing patterns of nonsynonymous and synonymous changes reject neutrality in favor of positive selection on various branches in the phylogeny after duplication (Figure 2, 4). Not only did the molecular clock rate increase for both passerine GHs (Figure 1; Supplemental Table 6), but the GH2 paralog experienced a burst of positive selection directly after the duplication event, driving its amino acid sequence apart from the ancestral one. Third, GH1 and GH2 retain key regulatory and structural sequence features necessary for the expression and function of growth hormone family proteins (Figures 4, 5, 6). The combination of large-scale structural conservation together with accelerated evolution at mostly differing amino acid sites in the two paralogs suggests that they may have evolved novel or specialized functions beyond simple dosage effects.

From the syntenic and sequence evidence we present, it is evident that the two copies of passerine GH can be explained by a single short segmental duplication and interchromosomal translocation in the common ancestor of Passeriformes. This mechanism contrasts with other expansions of the growth hormone family which have taken place primarily via tandem duplication (Papper et al. 2009; Wallis and Wallis 2006; Alam et al. 2006; Wallis et al. 1998) and whole genome duplication (Whittington and Wilson 2013). In passerines, post-duplication deletions and a series of rearrangements on chromosome 1 have altered the syntenic context and possibly the regulatory milieu of GH1 and GH2 among the major passerine lineages. In addition, the base composition of the paralogs has diverged substantially post-duplication. This may be a consequence of translocation onto a macrochromosome, as avian micro- and macrochromosomes differ on average in their GC content (Warren et al. 2017; O’Connor et al. 2019). GC content can influence the transcriptional state of chromatin (Tillo and Hughes 2009), and thus may contribute to divergence in expression patterns of the paralogs.

Our work, using the rich genomic resources now publicly available for birds, builds on the discovery of passerine GH2 and initial tests of evolutionary rate acceleration and positive selection performed by Yuri et al. (2008). Our data confirm their observation of a rate increase in amino acid sequence evolution of both passerine GH1 and GH2 in comparison to non-passerine avian GH. That Yuri et al. estimated larger rate increases than those we found (Supplemental Table 6) is likely due to their more limited taxon sample and use of partial sequences that happened to span a region (84 amino acids from sites 51-134) that is evolving more rapidly than the overall sequence (Figure 4). The rate increases in GH evolution we estimated from complete coding sequences of a much larger taxon sample are still distinctly higher than expected from any previously observed genome-wide rate changes. For example, Yuri et al. (2008) estimated a roughly 2X increase for passerine GH introns and Nam et al. (2010) estimated an average rate increase of 0.08X for coding sequences in the zebra finch genome by comparison of 8,384 genes with their 1: 1 orthologs from chicken. Yuri et al. (2008) detected evidence of positive selection on two amino acid sites, 99 and 107 as numbered herein. With a much larger taxon sample of complete sequences, we confirmed one of these sites (99) and found further evidence of positive selection on a total of 21 sites, 11 in each paralog, one of which is shared in GH1 and GH2.

The shared site is amino acid 27, the last residue in the predicted signal peptide of the vast majority of avian GH genes (Supplemental Table 3). Signal peptidase is predicted to cut these polypeptides between sites 27 and 28, so 27 is in the −1 position with respect to cleavage. The −1 position is crucial for correct recognition and cleavage by signal peptidase (Auclair et al. 2012). Mutations here may destroy the signal peptidase cleavage site altogether if the replacement residue is incompatible, such as one with a polar or bulky side chain. Other mutations may affect the efficiency of signal peptidase binding (Choo and Ranganathan 2008). Therefore, positive selection on site 27 indicates evolutionary pressure for the two passerine paralogs to interact differently with signal peptidase. Consistent with diverging roles for the two paralogs, 64 of 154 passerine species sampled have lost a predicted signal peptidase cleavage site in either GH1 or GH2, most often in GH2. In many other species, the probability that such a site is present is substantially reduced. Changes destroying or modifying signal peptidase’s affinity would likely affect whether GH proteins are exported from the cell, and if so, how efficiently. Export is, of course, a crucial step in the typical function of a protein hormone. GH genes without signal peptides may be specialized for intracellular roles of the protein.

Other positively selected sites are clustered at one end of the 3D model of the folded protein (Figure 5A). One of the disulfide bridges and site 90, which has the Gly>Lys replacement in all GH2, are also found in this region (Figure 5B). Disulfide bridges are often integral to maintaining protein tertiary structure, and site 90 is known to be involved in binding to the GH receptor in humans. The proximity of many positively selected sites to these features suggests that selection may be acting to further modify protein function by changing binding affinity to the GH receptor or other ligands.

Degeneration or silencing are the most common fates of duplicate genes. However, the presence of many universally conserved sites and features suggests that the amino acid sequences of these paralogs are not in the process of degeneration. Among these conserved features are the four core alpha helices, the universally conserved cysteines expected to form two disulfide bridges, one site for potential phosphorylation, and most regions of the mature protein that likely interact with the growth hormone receptor. Rather than degenerating, each paralog may be freed from some selective constraints and adapting under positive selection.

Neither have these paralogs been silenced. Instead, gene expression data reported here and in previous studies suggests divergent patterns of GH1 and GH2 expression depending on species, and, possibly, lineage. In the suboscine *Pipra*, GH2 expression was limited entirely to the pituitary among the tissues we examined, while GH1 was expressed in multiple tissues. In the oscine *Corvus*, GH2 was expressed in a wider range of extrapituitary tissues than GH1 (Arai and Iigo 2010). Both GH1 and GH2 are expressed in brain of the oscine zebra finch, but only GH1 is implicated in the song learning process. (Xie et al. 2010). Song learning is a defining trait of oscines that is thought to be ancestral in Passeriformes, but lost in many suboscine lineages (Odom et al. 2014). Differing patterns of GH paralog expression may be influenced by rearrangements of chromosome 1 affecting the region upstream of GH2, which have occurred in both oscines and suboscines. Base substitutions that alter key regulatory elements of the promoter regions immediately upstream of both paralogs (Figure 6) may also be involved in observed expression differences (Figure 7).

These divergent patterns of tissue-, species- and stimulus-specific patterns of gene expression are consistent with post-duplication evolution of the passerine GH paralogs to novel or more specialized roles through neofunctionalization or subfunctionalization. The pattern reported for the oscine *Corvus* is more consistent with neofunctionalization of GH2, as GH2 is expressed in novel tissue types compared to ancestral GH1 (Arai and Iigo 2010). In contrast, the pattern we report in the suboscine *Pipra* is more consistent with subfunctionalization of GH2, with GH1 being found across tissues matching ancestral growth hormone expression observed in chicken (Harvey and Hull 2003; Martínez-Moreno et al. 2014), but with GH2 limited to the pituitary. One possibility is that the pituitary functions of ancestral GH1 have been subdivided in passerine GH1 and GH2.

The duplication of a gene so crucial to life history strategy, body composition, and reproduction may be a key adaptation driving the acceleration of passerine development, metabolism, and shorter generation times. These factors may also help explain a faster mutation rate (Gillooly et al. 2005; Nabholz et al. 2008), which correlates to increased diversification rates in birds (Lanfear et al. 2010). This accelerated evolution may have aided adaptation to novel environments as the passerine radiation spread across the globe from its putative origin on the Australian landmass (Oliveros et al. 2019; Barker et al. 2004).

Further research is needed into the developmental stage- and tissue-specific expression patterns of the two paralogs. This fundamental biological data will help elucidate the roles of these duplicate genes in passerine growth and development. Our investigations of the molecular evolution and gene expression of the passerine GH paralogs lend powerful evidence to the adaptive fates of these GH paralogs post-duplication. Our findings show the retention of function and expression of both passerine GH paralogs, and suggest that they are evolving in a manner consistent with novel or specialized functions. These derived functions may include crucial adaptations to the accelerated life history strategy associated with passerine diversity.

## Materials and Methods

### Determining copy number and synteny

For our initial descriptive analysis, we used 76 well-annotated avian genomes (24 passerine, 52 non-passerine) available on NCBI (O’Leary et al. 2016), plus the unannotated chromosome-level genome assemblies for the passerines *Acanthisitta chloris* and *Manacus candei*. Source genomes used in these and following analyses are listed in Supplemental Table 1. To identify GH sequences and copy number, we searched annotated genomes for “somatotropin” and “growth hormone,” which yielded hits for genes annotated “somatotropin” “somatotropin-like” “growth hormone” “GH” and “GH1.” We then used the mRNA sequence of each annotated gene to query the focal taxon (taxid matching the genome of origin) using blastn. We evaluated the top 10 matching hits (or all hits, if fewer than 10 resulted). From this pool, we retained all hits matching genomic sequences with a sequence identity above 60%, a gapless coding sequence (CDS), and no annotation that better represented the identity of the matching region.

We examined the syntenic context of these annotated growth hormone genes. When chromosome-level assemblies were available, we noted the chromosomal location of the GH paralog(s). For each annotated avian genome, we recorded the genes flanking GH genes. For non-passerines, we also noted the two genes flanking either side of the homologous genomic region where a segment containing GH2 is inserted in passerines (Figure 2). Additionally, for the acanthisittine and suboscine syntenies, we used blast, querying genes annotated on same species’ or close relatives’ genomes, to identify genes on unannotated chromosome-level assemblies for *Manacus candei* and *Acanthisitta chloris*. Many annotated genomes did not convey enough information to reconstruct the full syntenies shown in Figure 2 due to scaffolds ending in the middle of key regions. For a list of genomes informing the placement of the genes in these regions, see Supplemental Table 8.

To characterize changes that have occurred between passerine lineages in the local genomic context of GH2, we used the chromosomal and local syntenic context of GH genes as well as pairwise dotplots to visualize runs of high sequence similarity. We extracted sequences 25 Mb in length centered on the GH2 translation start site from chromosome-level genome assemblies from each passerine suborder: oscine (*Taeniopygia guttata*), suboscine (*Chiroxiphia lanceolata*), acanthisittine (*Acanthisitta chloris*). To obtain dotplots focused on the displacement of the segment containing GH2 in oscines compared to suboscines and *Acanthisitta*, we ran pairwise alignments of these three sequences in D-GENIES (Cabanettes et al. 2018) using the Minimap2.24 aligner (Li 2018), selecting “many repeats”, and mapping results with the R version 4.2.0 (R Core Team 2022) package pafr (Winter 2020)(Supplemental Figure 1).

### Growth hormone sequence curation and alignment

For the initial categorization of GH paralogs as GH1 (ancestral) or GH2 (duplicate), we used sequences from the set of 76 annotated avian genomes from NCBI (O’Leary et al. 2016). GH paralogs were categorized as ancestral (GH1), or novel (GH2), based on their annotated syntenic context. To retrieve additional GH sequences, we queried 280 unannotated genomes with BLAST using the GH sequence(s) of the closest available annotated relative (Supplemental Table 1). We assigned retrieved unannotated sequences the identity of GH1 or GH2 based on sequence similarity to annotated GH1 and GH2 paralogs. Retrieved sequences underwent quality checks: complete coding DNA sequences without multiple stop codons, gaps, or Ns were retained, as were sequences with other nucleotide ambiguities (R, Y, S, W, K, M). Failure to retrieve one full sequence from certain non-passerine genomes or two full sequences from certain passerine genomes was most likely due to genome incompleteness or mis-assembly. One sequence (*Struthio camelus* GH1) was discarded because it was highly divergent from all other bird sequences and contained multiple stop codons, also likely due to mis-assembly. We calculated base composition on our dataset of avian GH sequences that were free of ambiguities.

For signal peptide prediction, we used amino acid translations of the same avian GH sequences as used in the selection tests described in Methods: *Tests of Selection*. For each of these sequences, we used SignalP 6.0 (Teufel et al. 2022) to predict the likelihood of each sequence containing a signal peptide. For sequences where the likelihood was > 0.5, SignalP 6.0 predicted both the presence of a cleavage site and the probability of the cleavage site being in the predicted location. We chose the settings “Eukarya” to limit signal peptide prediction sites to Sec/SPI sites only, as well as the “slow” setting to use the more complex and accurate protein language model (Teufel et al. 2022).

We performed the coding CDS alignment (Supplemental Data: GH CDS Alignment) in Geneious (Geneious® 2020.2.5) using Clustal Omega and then visually checked it. To analyze the upstream cis-regulatory region, we retrieved 1 kb sequences 5’ of the translational start site, plus the first three codons to anchor the alignment, from 132 passerine GH1s, 146 GH2s, and five non-passerine avian GHs. We aligned these 1009 bp sequences with the Geneious plugin of MAFFT v. 7 (Katoh and Standley 2013) using default settings: algorithm = auto, gap open penalty 1.53, offset value 0.123, scoring matrix 200 PAM/k = 2. We then annotated the 1009 bp chicken sequence with the regulatory elements identified by Ip et al. (2004), and realigned the 250 bp region proximal to the translation start site with the same default settings in MAFFT to increase the accuracy of the alignment around regulatory element binding sites (Supplemental Data: GH Flanking Region Alignment), as the full 1009 bp alignment featured numerous indels affecting the overall alignment. We considered sequence motifs aligning with the regulatory elements annotated on the chicken sequence to be homologous after inspection of the alignment.

### Nomenclature

Given the clear evolutionary origin to the GH paralogs we found (see Results: *Copy number and duplication origin*), for passerine sequences, we chose to use the symbol GH1 to refer to the ancestral paralog and GH2 to refer to the novel derived paralog. GH is used to refer to the non-passerine avian paralog as per the Chicken Gene Nomenclature Consortium (Burt et al. 2009). Previous authors, who had less data from fewer species, have used a variety of nomenclature. Yuri et al. (2008) named the paralogs based on the PCR product length of the partial passerine sequences examined, with GHL (Growth Hormone Long) having a longer intron than GHS (Growth Hormone Short). However, gene lengths are variable and some ancestral paralog sequences are shorter than novel paralog sequences (e.g. *Catharus ustulatus* has a 4166 bp GH1 and a 5033 GH2; *Passer domesticus* has a 4287 bp GH1 and 5206 bp GH2). Arai and Iigo (2010) assigned the novel copy the name GHA and the ancestral copy GHB; but these turned out to be unintuitive given the evolutionary history now evident. Xie et al. (2010) reference GH1 (ancestral) and GH1A (novel); while workable, we argue that GH1 and GH2 are clearest.

### Phylogenetic Tree Estimation

We estimated the trees used in the evolutionary analyses from the Geneious alignment of GH CDS using RAxML v. 8 (Stamatakis 2014) option “-f a,” which searches for the best maximum likelihood scoring tree and performs a rapid bootstrap analysis in one program run. We used 1000 bootstrap replicates under the GTRCAT model. Two topologies were obtained and used in all evolutionary analyses: one from an unconstrained maximum likelihood tree search (the unconstrained tree) and a second (the constrained tree) in which the search was constrained to trees compatible with an input topology specifying many widely accepted relationships of the taxa involved (See Supplemental Figure 2 for the unconstrained and constrained topologies, as well as the input constraint topology). Many of the constrained relationships represent short internodes deep in the tree; such relationships are unlikely to be recovered with a short single gene CDS alignment but are well supported by much larger phylogenomic datasets. Both trees were used throughout all evolutionary analyses to test for any effect of using the constrained tree versus unconstrained tree, making the biologically plausible assumption that the actual evolutionary history of GH genes (aside from duplication in the passerine clade) may be better represented by the constrained topology formed from literature consensus derived from a multitude of sites. Relationships still unresolved in the avian literature were not constrained (e.g. the placement of *Opisthocomus hoazin* within Neoaves). The constraint topology was invoked in RAxML with the -g setting.

### Molecular clock

We performed tests of the relative molecular clock rate of protein-coding nucleotide sequence evolution on the constrained and unconstrained trees with the BASEML package in PAML version 4.9j (Yang 2007). The null model enforced a global clock rate, while alternative models allowed local clock rates to be determined for designated foreground clades. For both phylogenetic topologies tested, we designated each of the following as the foreground clade in a separate test: passerine GH1, passerine GH2, and passerine GH1 + GH2. We also performed a fourth test with each topology in which passerine GH1 and GH2 were allowed to have separate clock rates. See Figure 1 for a visualization of these foreground and background branch sets.

### Tests of Selection

We estimated the timing and sequence location of positive selection in the avian GHs with branch-site tests. We used the codeml program in PAML v. 4.9j (Yang 2007) to estimate the branch- and site-specific ratios of nonsynonymous to synonymous substitutions (dN/dS or ω). In the branch-site test, the null model M7 (all branches and sites had a beta-distributed ω < 1) is compared to various alternate M8 models. M8 specifies the M7 condition on background branches and sites, allowing an additional category of ω on user-specified foreground branches (including all of their sites) where ω > 1, indicating elevated nonsynonymous changes and positive selection on branches and sites where this rate category occurred. We specified the following foreground clades: A) the passerine GH1 clade, B) the passerine GH2 clade, C) all passerine GH (both GH1 and GH2), and D) the two branches immediately following the duplication event to explicitly test the hypothesis of a burst of positive selection following duplication (Figure 1). The tests on A, B, and C are discussed in Results, but the test on clade D failed to estimate ω for this test, likely due to the small number of test branches compared to the size of the dataset, so it was discarded. We also employed the aBSREL test (Smith et al. 2015) in the HyPhy 2.5 family of methods (Kosakovsky Pond et al. 2020) using the Datamonkey server (Weaver et al. 2018). aBSREL represents a stricter version of the codeml branch-site test, as it allows more rate categories of ω, including ω > 1 on background branches. We tested the same A, B, C, and D foreground branch sets as in codeml.

We then employed the Bayes Empirical Bayes test in the codeml program on the GH1 and GH2 clades individually to identify amino acids under selection. We placed the amino acids identified under positive selection on the translated CDS alignment, along with other amino acid features identified as important in the growth hormone literature (see Figure 4), as well as on a 3D structural model of *Gallus gallus* GH available from AlphaFold v. 2.1.1 (Jumper et al. 2021, Varadi et al. 2021) (Figure 5). We also modeled zebra finch GH1 and GH2 sequences with AlphaFold v. 2.1.1 to confirm that major structural features (alpha helices, disulfide bridges) had the same predicted placement as in chicken GH (data not shown). To capture paralog sequence differences that may not show up as positively selected sites, we visually examined the amino acid alignment of avian GH, GH1, and GH2 for fixed mutations between either passerine paralog and all other sequences.

### Characterization of Expression of GH Axis Genes with RNA-seq

As part of a separate study on male cooperative display behavior (Horton et al. 2019), twelve male wire-tailed manakins (*Pipra filicauda*) were collected from the wild and sacrificed at Tiputini Biodiversity Station, Ecuador. Whole brains, pituitary, and testes were extracted from birds 4-6 minutes postmortem and stored in dry ice until placed in liquid nitrogen.

The brains were cryosectioned and microdissected for 10 individual brain nuclei known to be relevant to social behavior and sex-hormone regulation: the Ventromedial Hypothalamus (VMH), Anterior Hypothalamus (AH), Paraventricular Nucleus (PVN), Medial Preoptic Area (POM), Dorsomedial Intercollicular Nucleus (ICO), Midbrain Central Gray (GCT); part of the Mesolimbic Reward System: Arcopallium Intermedium (AI); or part of both: Bed Nucleus of the Stria Terminalis (BSTm), Lateral Septum (LS), Nucleus Taenia (TNA). The mRNA from these ten nuclei, pituitary gland and testis, were extracted using a Qiagen miRNeasy Micro Kit (Bolton PE, unpublished data; Horton et al. 2019).

The ten brain nuclei and two peripheral tissues were prepared as TruSeq Stranded mRNA Sample Prep Kit libraries per tissue and per individual. Sequencing was performed on an Illumina HiSeq 4000. We mapped resultant single-ended reads to the annotated *Pipra filicauda* genome (GCA_003945595.1) using splice-aware mapper STAR v. 2.75 (Dobin et al. 2013) on two pass mode. We set mismatch tolerance to 0.04, allowing four mismatches per 100 bp read, which is less than the sequence divergence of each exon between *Pipra* GH1 and GH2, except the very short exon 1. Visual inspection in IGV showed no bias for mapping rates in exon 1 of either paralog. We calculated the total number of reads per gene and per exon using featureCounts v2.0.1 (Liao et al. 2014). We used a counts-per-million reads (CPM) formula to calculate a length normalized count for genes using the total exon length. For detailed methods and permits for animal collection, import, and export, see Horton et al. (2019).

## Supporting information

Supplemental Data: Amino Acid Alignment

Supplemental Data: GH CDS Alignment

Supplemental Data: Flanking Region Alignment

## Acknowledgements

We are grateful to Josefin Stiller, Shaohong Feng, Guojie Zhang, Erich Jarvis, and Olivier Fedrigo for granting access and early results on the B10K round 2 genomes. We thank Karen Carleton, Carlos Machado, Tom E. Porter, Ed Braun, Rebecca Kimball, and Barney Schlinger for their insightful feedback on this work; we also thank Alexey Zimin, Rebecca Dikow and Paul Frandsen for assembling the red siskin genome; lastly, we thank Chris Balakrishnan, Brent Horton, Brandt Ryder, Ros Dakin, Ignacio Moore and Jennifer Houtz for collecting and dissecting tissues in the field for the *Pipra filicauda* transcriptome dataset.

## Notes

### Competing Interest Statement

The authors have declared no competing interest.

