## Supplemental Figure 1 for "Evolution of the Growth Hormone Gene Duplication in Passerine Birds"

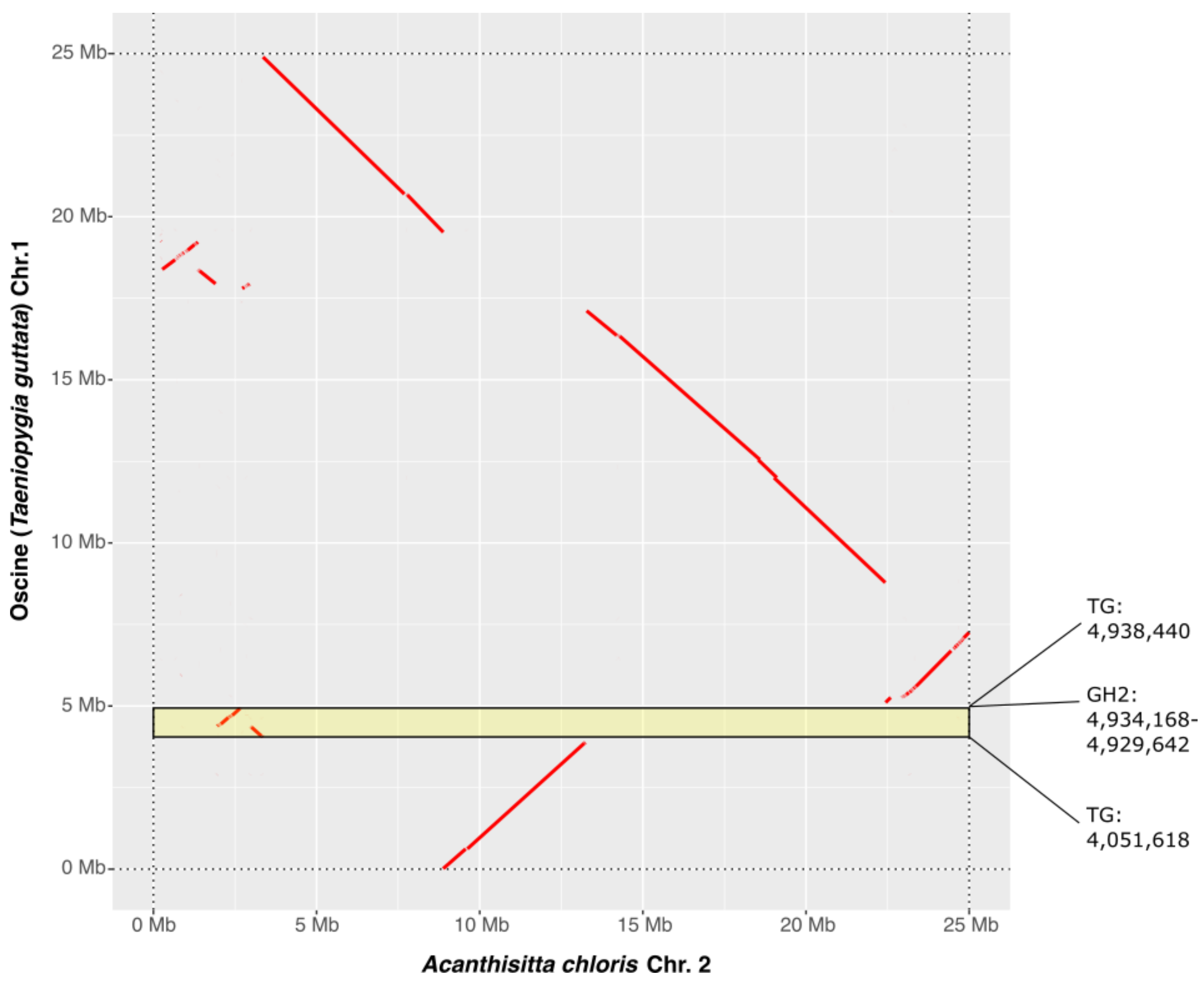


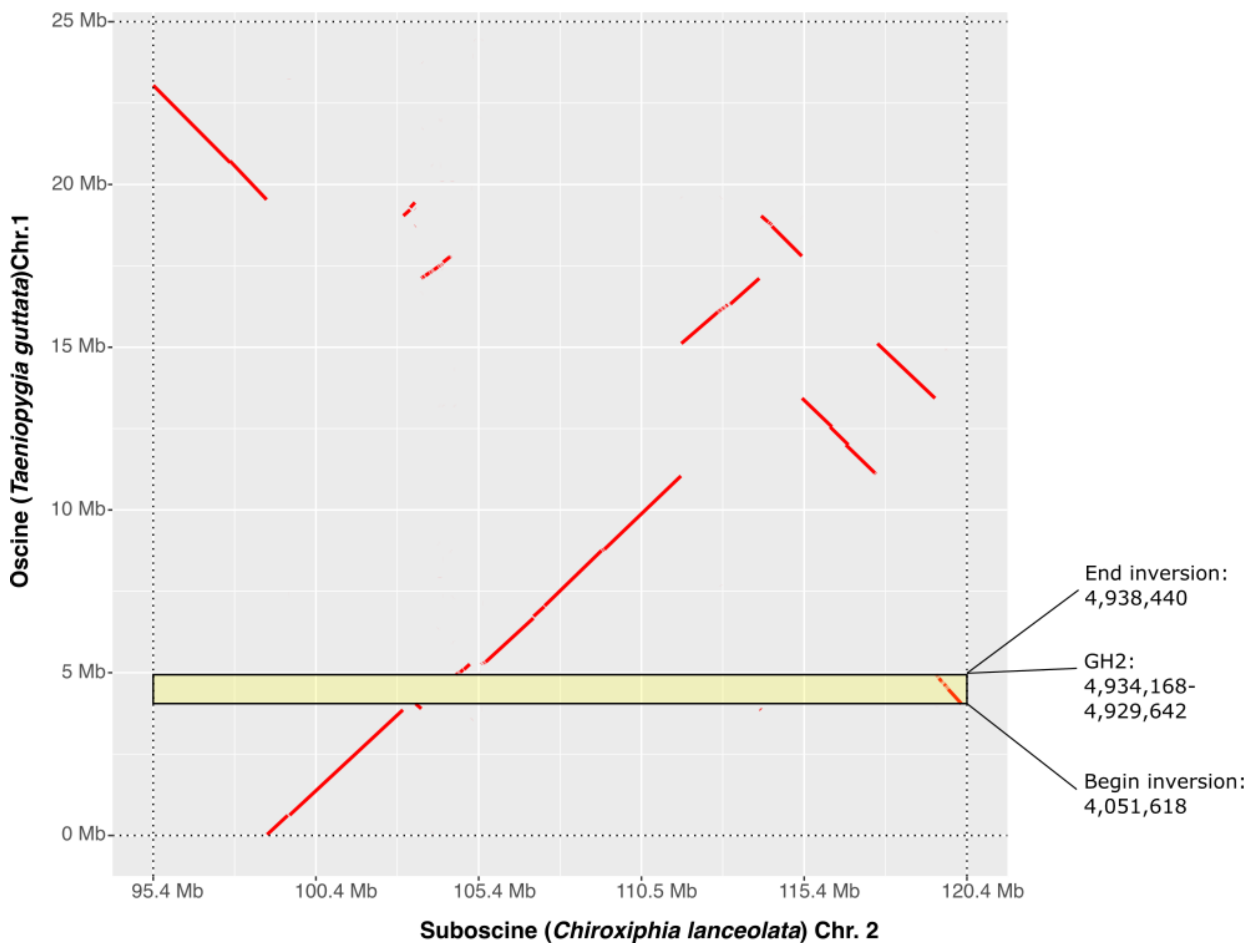


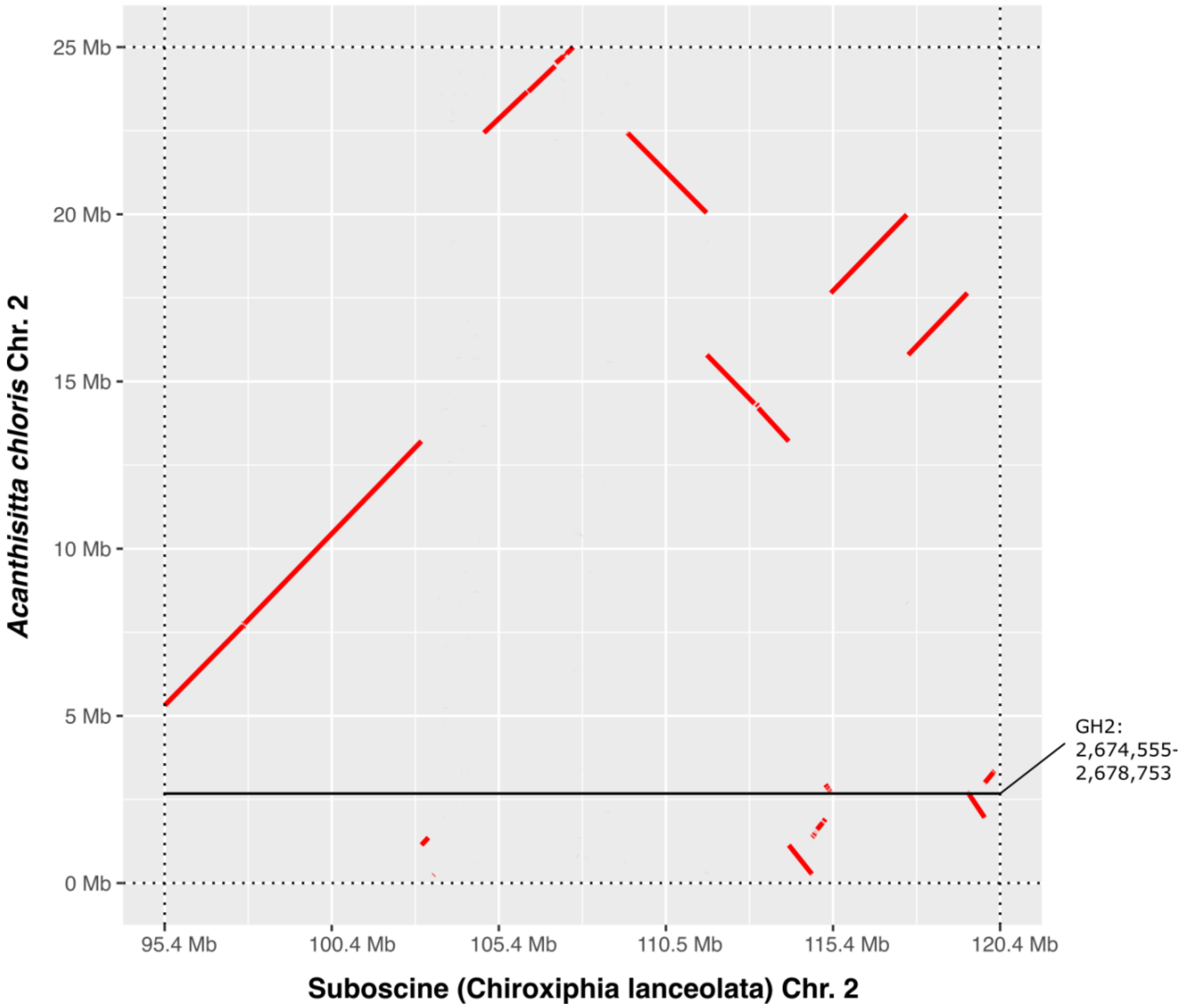


***Supplemental Figure 1: Top left panel:*** *the first 25 megabases (Mb) and final 25 Mb of the oscine chromosome 1 and the homologous acanthisittine chromosome 2 are aligned and displayed in a dotplot. GH2 is on an inverted and displaced segment. The beginning and ending coordinates of the inversion in* Taeniopygia guttata *estimated in the alignment are labeled, with the coordinates for GH2 listed also. The inversion breakpoint is an estimated 4 kb from the GH2 translation start site.* ***Top right panel:*** *the first 25 Mb of the oscine chromosome 1 and the homologous suboscine chromosome 2 are aligned and displayed in a dotplot. GH2 is on a displaced segment. The highlighted box is for the same coordinates of* Taeniopygia guttata *chr. 1 in both top panels.* ***Bottom left panel:*** *the first 25 Mb of the acanthisittine chromosome 2 and the final 25 Mb of the suboscine chromosome 2 are aligned. The location of GH2 on* Acanthisitta chloris *chr. 2 is marked.*
