## Supplemental Figure 2 for "Evolution of the Growth Hormone Gene Duplication in Passerine Birds"

**Supplemental Figure 2:** Tree Topologies used in tests of evolutionary rates and selection. **A.** The unconstrained gene tree topology obtained with RAxML using the growth hormone coding sequence alignment (Supplemental Data: GH CDS Alignment). **B.** The constrained topology obtained using the same CDS alignment as in A, but with the tree search constrained to trees compatible with an input topology. **C.** The input constraint topology used to obtain B. The input topology captures well-resolved and widely-accepted species relationships of birds based on prior studies, a fraction of which are cited below. RAxML settings used for A and B are in Methods: *Phylogenetic Tree Estimation*.

**A. Unconstrained Tree Topology**

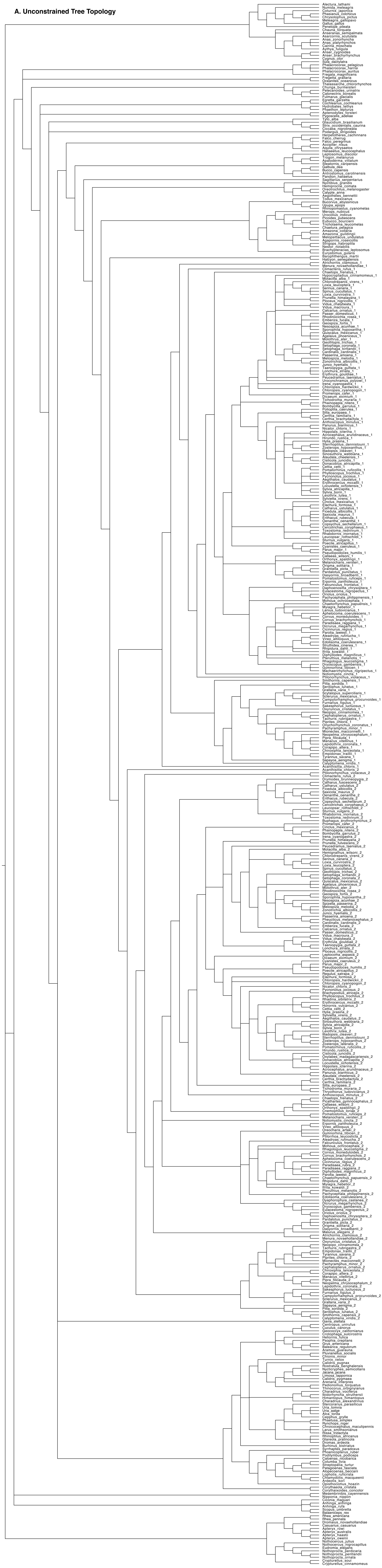

**B. Constrained Tree Topology**

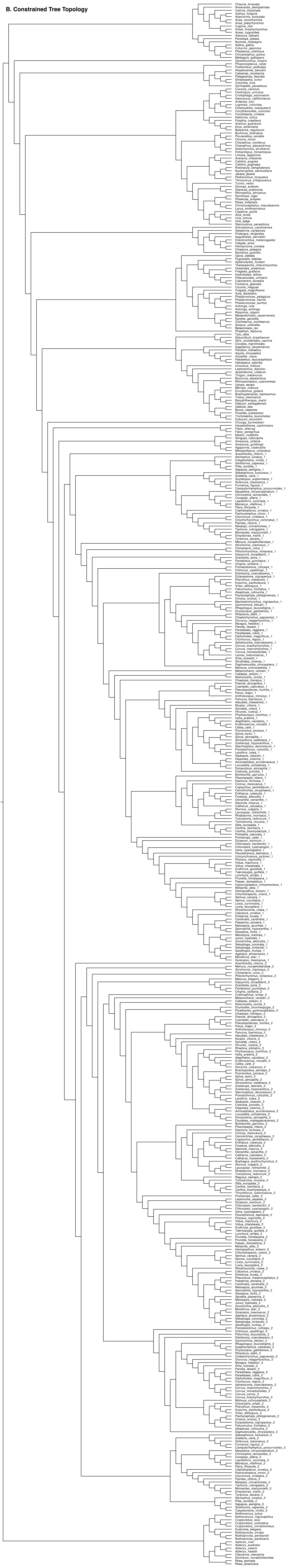

**C. Constrained Tree Backbone**

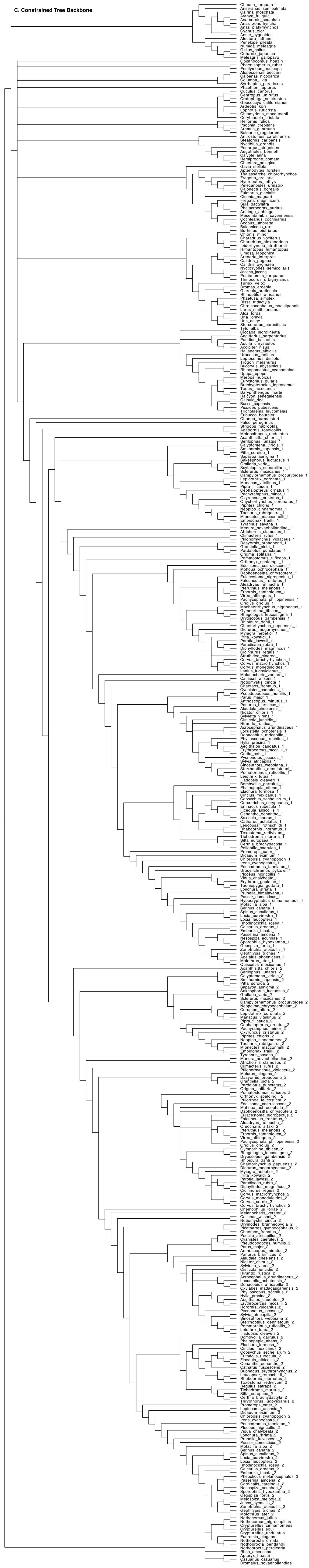
