## Supplemental Figure 3 for "Evolution of the Growth Hormone Gene Duplication in Passerine Birds"

### Supplemental Figure 3:
### Phylogenetic Distribution of 15 bp Repeat in GH1 Promoter Region

**Legend:**
- 15 bp tandem repeat absent
- 15 bp tandem repeat present
- no data

**Supplemental Figure 3. The phylogenetic distribution of the 15 bp tandem repeat is shown on the "Constrained Tree" topology (see Methods: Phylogenetic Tree Estimation; Supplemental Figure 2). As this repeat is only present in the promoter region of passerine GH1, the passerine GH1 clade is shown here. Red leaf labels indicate one copy of the 15 bp motif in that species; green leaf labels indicate two copies (see Supplemental Data: GH Flanking Region Alignment). Black leaf labels indicate that no promoter region was retrieved for this species' GH1. Internal branches are colored red or green based on simple parsimony.**

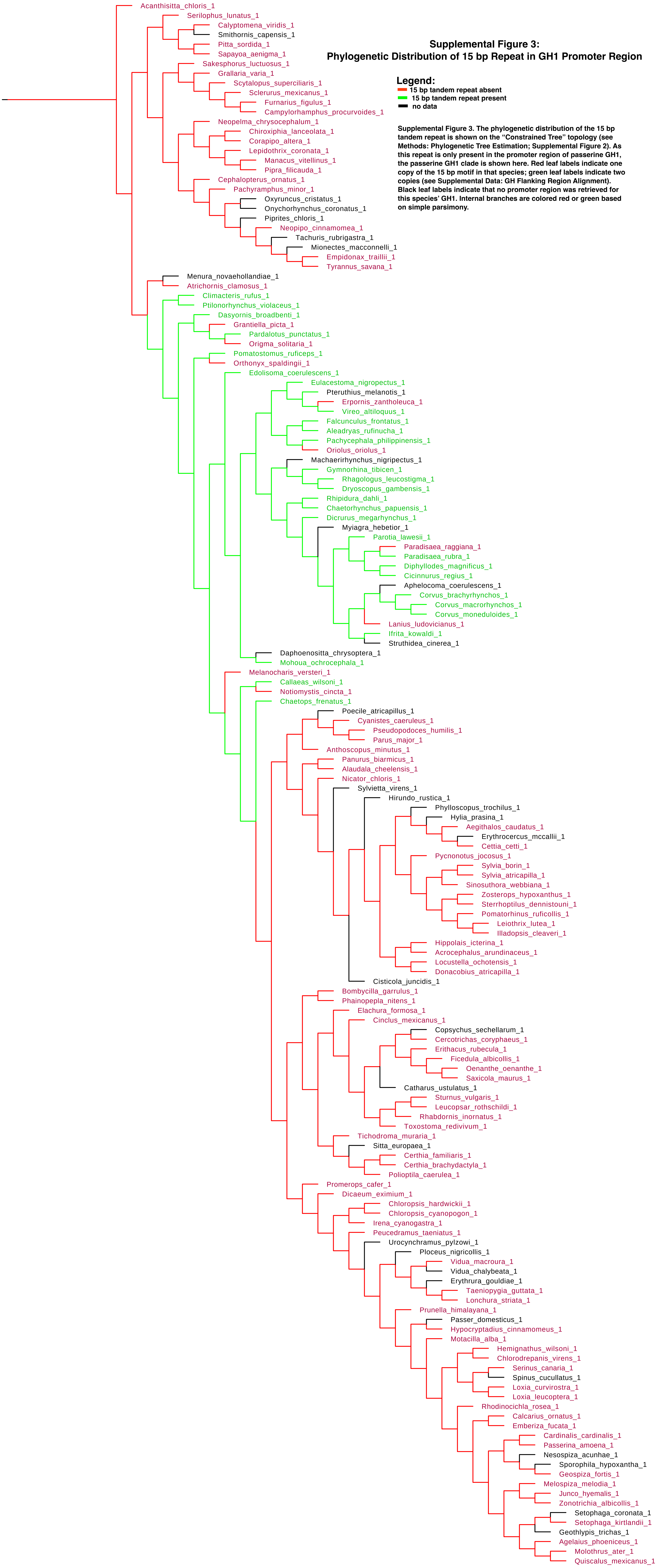
