## Supplemental Table 4 for "Evolution of the Growth Hormone Gene Duplication in Passerine Birds"

| Paralog | 5’ UTR Length | 3’ UTR Length | Suborder |
|---|---|---|---|
| **Passerine GH1** | | | |
| *Acanthisitta chloris* GH1 | 49 | 36 | Acanthisitta |
| *Corapipo altera* GH1 | 60 | 47 | Suboscine |
| *Empidonax trailli* GH1 | 60 | 71 | Suboscine |
| *Manacus vitellinus* G1 | 47 | 47 | Suboscine |
| *Neopelma chrysocephalum* GH1 | 54 | 22 | Suboscine |
| *Corvus brachyrhynchos* GH1 | 97 | 53 | Oscine |
| *Corvus moneduloides* GH1 | 97 | 53 | Oscine |
| *Cyanistes caeruleus* GH1 | 25 | 50 | Oscine |
| *Geospiza fortis* GH1 | 36 | 51 | Oscine |
| *Lonchura striata* GH1 | 41 | 51 | Oscine |
| *Parus major* GH1 | 25 | 50 | Oscine |
| *Pseudopodoces humilis* GH1 | 40 | 50 | Oscine |
| *Serinus canaria* GH1 | 40 | 51 | Oscine |
| *Sturnus vulgaris* GH1 | 21 | 51 | Oscine |
| *Zonotrichia albicollis* GH1 | 40 | 51 | Oscine |
| **Average Oscinene GH1** | **48.8** | **48.93333333** | |
| **Passerine GH2** | | | |
| *Acanthisitta chloris* GH2 | 65 | 96 | Acanthisitta |
| *Neopelma chrysocephalum* GH2 | 61 | 100 | Suboscine |
| *Corvus brachyrhynchos* GH2 | 77 | 96 | Oscine |
| *Corvus cornix* GH2 | 76 | 96 | Oscine |
| *Corvus moneduloides* GH2 | 77 | 96 | Oscine |
| *Geospiza fortis* GH2 | 59 | 91 | Oscine |
| *Lonchura striata* GH2 | 61 | 96 | Oscine |
| *Parus major* GH2 | 60 | 91 | Oscine |
| *Pseudopodoces humilis* GH2 | 60 | 98 | Oscine |
| *Serinus canaria* GH2 | 101 | 98 | Oscine |
| *Sturnus vulgaris* GH2 | 61 | 87 | Oscine |
| *Taeniopygia guttata* GH2 | Not Annotated | 83 | Oscine |
| *Zonotrichia albicollis* GH2 | 59 | 89 | Oscine |
| **Average Oscinene GH2** | **68.08333333** | **93.61538462** | |
| **Non-passerine Avian GH** | | | |
| *Apaloderma vittatum* GH | 58 | 21 | |
| *Aptenodytes forsteri* GH | 52 | 90 | |
| *Apteryx mantelli* GH | 58 | 145 | |
| *Cariama cristata* GH | 58 | 95 | |
| *Chaetura pelagica* GH | 45 | 97 | |
| *Charadrius vociferus* GH | 58 | 94 | |
| *Cuculus canorus* GH | 53 | 95 | |
| *Fulmarus glacialis* GH | 28 | 90 | |
| *Gallus gallus* GH | 33 | 97 | |
| *Gavia stellata* GH | 58 | 112 | |
| *Merops nubicus* GH | 58 | 18 | |
| *Nothoprocta perdicaria* GH | 119 | 82 | |
| **Average Non-passerine Avian GH** | **56.5** | **86.33333333** | |
| *Homo sapiens* GH1 | 63 | 106 | |

**Supplemental Table 4: Length of Untranslated Regions (UTRs)** Table of 5’ and 3’ Untranslated Region (UTR) lengths retrieved from NCBI annotated GH genes available on NCBI Genome Data Viewer (Rangwala et al. 2020). Relatively few avian GH genes have UTRs annotated. In addition to the 5’ and 3’ UTR length listed per species and paralog, the mean length per avian paralog (GH1, GH2, GH) is given. For passerine species, the suborder is listed. At the bottom of the table, the well-characterized human GH1 UTRs are given for comparison.
