## Supplemental Table 6 for "Evolution of the Growth Hormone Gene Duplication in Passerine Birds"

| Input Tree Topology | Model | Likelihood (lnL) | Rate for GH1 | Rate for GH2 | Tree Length | P-value |
|---|---|---|---|---|---|---|
| Gene Only | H0 | -28817.61181 | 1 | 1 | 13.55847 | |
| Gene Only | H1 | -28787.38673 | 2.03408 | 1 | 11.67174 | <.00001 |
| Gene Only | H2 | -28781.49987 | 1 | 1.96101 | 11.06946 | <.00001 |
| Gene Only | H3 | -28715.06552 | 5.73077 | 5.73077 | 6.31309 | <.00001 |
| Gene Only | H4 | -28714.8123 | 5.92727 | 5.45819 | 6.33981 | <.00001 |
| Constrained | H0 | -29173.86902 | 1 | 1 | 15.05307 | |
| Constrained | H1 | -29149.28404 | 1.8924 | 1 | 13.10197 | <.00001 |
| Constrained | H2 | -29152.52407 | 1 | 1.70865 | 12.71838 | <.00001 |
| Constrained | H3 | -29099.04514 | 4.59198 | 4.59198 | 7.41313 | <.00001 |
| Constrained | H4 | -29098.66611 | 4.79045 | 4.28941 | 7.4615 | <.00001 |

**Supplemental Table 6A:** Relative rate tests in baseml of coding sequence evolution comparing the passerine clades (GH1, GH2) to non-passerine avian GH clades. The input tree topologies are "Gene Only," the tree estimated only based on GH coding sequences, and "Constrained," which is a GH gene tree constrained to be compatible with an input tree based on widely accepted avian species relationships (see Methods and Supplemental Figure 2). The null model, H0, tests a single rate on the whole GH tree. The passerine GHs are allowed different rates in the alternate models: H1 allows a different rate for the GH1 clade; H2, the GH2 clade; H3, a different rate for GH1+GH2; H4, three rates, meaning GH1, GH2 are allowed to vary compared to the background. The tree length refers to the sum of branch lengths in each estimate based on the expected number of substitutions. The null model was compared to each alternate model, with the p-value shown.

| Input Tree Topology | Model | Likelihood (lnL) | Test Statistic | P-value |
|---|---|---|---|---|
| Gene Only | H0 | -28839.70818 | - | - |
| Gene Only | H1 | -28766.914211 | 145.58594 | <.00001 |
| Gene Only | H2 | -28679.567511 | 320.28134 | <.00001 |
| Gene Only | H3 | -28661.23946 | 356.93744 | <.00001 |
| Constrained | H0 | -29456.519124 | - | - |
| Constrained | H1 | -29358.444986 | 196.14826 | <.00001 |
| Constrained | H2 | -29301.435438 | 310.16736 | <.00001 |
| Constrained | H3 | -29251.618188 | 409.80186 | <.00001 |

**Supplemental Table 6B:** Branch-site tests for positive selection in codeml. The input tree topologies (Gene Only, Constrained) and clades tested in each model are the same as 2A. The null model, H0, does not allow for dN/dS > 1 (positive selection) on any site or branch. H1 allows for dN/dS > 1 in the passerine GH1 clade; H2 allows for dN/dS > 1 in the passerine GH2 clade, and H3 allows for dN/dS > 1 in the clade comprised of passerine GH1 and GH2. The test statistic is the likelihood ratio test between the null model and each alternate model. The p-value is shown with 1 degree of freedom, as each alternate model has one additional parameter (the site class of dN/dS >1) compared to the nested null model.

| Site | GH1 | GH2 |
|---|---|---|
| 18 | 0.907 | 0.5 |
| 22 | 0.994* | 0.785 |
| 27 | 1* | 1* |
| 31 | 0.5 | 0.613 |
| 38 | 0.5 | 0.983* |
| 67 | 0.5 | 0.692 |
| 71 | 0.758 | 0.5 |
| 72 | 0.999* | - |
| 77 | - | 0.611 |
| 85 | - | 0.959* |
| 88 | - | 1* |
| 96 | 0.997* | - |
| 99 | - | 1* |
| 120 | 1* | - |
| 122 | - | 0.997* |
| 128 | 0.847 | - |
| 146 | 0.979* | - |
| 148 | - | 0.946 |
| 150 | 0.809 | - |
| 151 | 1* | 0.571 |
| 154 | 1* | - |
| 156 | - | 1* |
| 158 | - | 1* |
| 159 | - | 1* |
| 161 | 0.99* | - |
| 162 | 1* | - |
| 163 | 0.943 | - |
| 167 | - | 1* |
| 177 | 1* | - |
| 178 | - | 0.592 |
| 203 | - | 0.908 |
| 209 | - | 0.999* |

**Supplemental Table 6C:** Amino acids identified as likely under positive selection by the Bayes Empirical Bayes test. PAML output for the test lists every amino acid site that is p > 0.5 likely to be under positive selection; sites not listed did not appear in the output and are unlikely to be under positive selection in either passerine GH1 or GH2. Sites of p > 0.95 are marked with an asterisk.
